# PROTOSPACEJAM: AN OPEN-SOURCE, CUSTOMIZABLE AND WEB-ACCESSIBLE DESIGN PLATFORM FOR CRISPR/CAS9 INSERTIONAL KNOCK-IN

**DOI:** 10.1101/2023.10.04.560793

**Authors:** Duo Peng, Madhuri Vangipuram, Joan Wong, Manuel D. Leonetti

## Abstract

CRISPR/Cas9-mediated knock-in of DNA sequences enables precise genome engineering for research and therapeutic applications. However, designing effective guide RNAs (gRNAs) and homology-directed repair (HDR) donors remains a bottleneck. Here, we present protoSpaceJAM, an open-source algorithm to automate and optimize gRNA and HDR donor design for CRISPR/Cas9 insertional knock-in experiments. protoSpaceJAM utilizes biological rules to rank gRNAs based on specificity, distance to insertion site, and position relative to regulatory regions. protoSpaceJAM can introduce recoding mutations (silent mutations and mutations in non-coding sequences) in HDR donors to prevent re-cutting and increase knock-in efficiency. Users can customize parameters and design double-stranded or single-stranded donors. We validated protoSpaceJAM’s design rules by demonstrating increased knock-in efficiency with recoding mutations and optimal strand selection for single-stranded donors. An additional module enables the design of genotyping primers for next-generation sequencing of edited alleles. Overall, protoSpaceJAM streamlines and optimizes CRISPR knock-in experimental design in a flexible and modular manner to benefit diverse research and therapeutic applications. protoSpaceJAM is available open-source as an interactive web tool at protospacejam.czbiohub.org or as a standalone Python package at github.com/czbiohub-sf/protoSpaceJAM.

## INTRODUCTION

The development of gene editing technologies, fueled by the discovery of CRISPR/Cas systems, has transformed our ability to manipulate genomes. A key goal of gene editing is to add new features to the genome via the controllable insertion (“knock-in”) of functional payloads. Applications of CRISPR knock-in span both research and clinical programs, including the systematic characterization of gene function using fluorescent protein tags (1, 2), the introduction of genetic variants for the elucidation of disease mechanisms (3), or the integration of chimeric antigen receptor payloads in T cells (CAR-T) for immunotherapy (4). Reflecting the breadth of these applications, many CRISPR/Cas-mediated approaches have been developed to enable knock-in in a variety of contexts (reviewed in (5)). For example, homology-independent methods that rely on non-homologous end-joining (NHEJ) enable knock-in in non-dividing cells (6), while prime editing allows knock-in in the absence of double-strand breaks (7). The most common experimental approach for site-specific knock-in leverages homology-directed repair (HDR) (8). In HDR-based knock-in, a site-specific guide RNA (gRNA) is used to recruit CRISPR/Cas at a genomic target and induce a double-strand break, while an exogenous DNA sequence (“donor”) is provided to direct the integration of the desired payload. In the donor, the payload is flanked by sequences homologous to the target site (“homology arms”) that can template DNA repair by co-opting endogenous repair pathways (Figure 1A). Because HDR is controllable and precise, it remains the most widely used method for genomic knock-in, although its use is restricted to dividing cells (8). Technical advances are rapidly increasing the efficiency of HDR-based knock-in in key cell types such as human stem cells (9–11) or primary T cells (12). In addition, CRISPR/Cas knock-in is now entering the clinic for CAR-T therapy (4).

**Figure 1.**
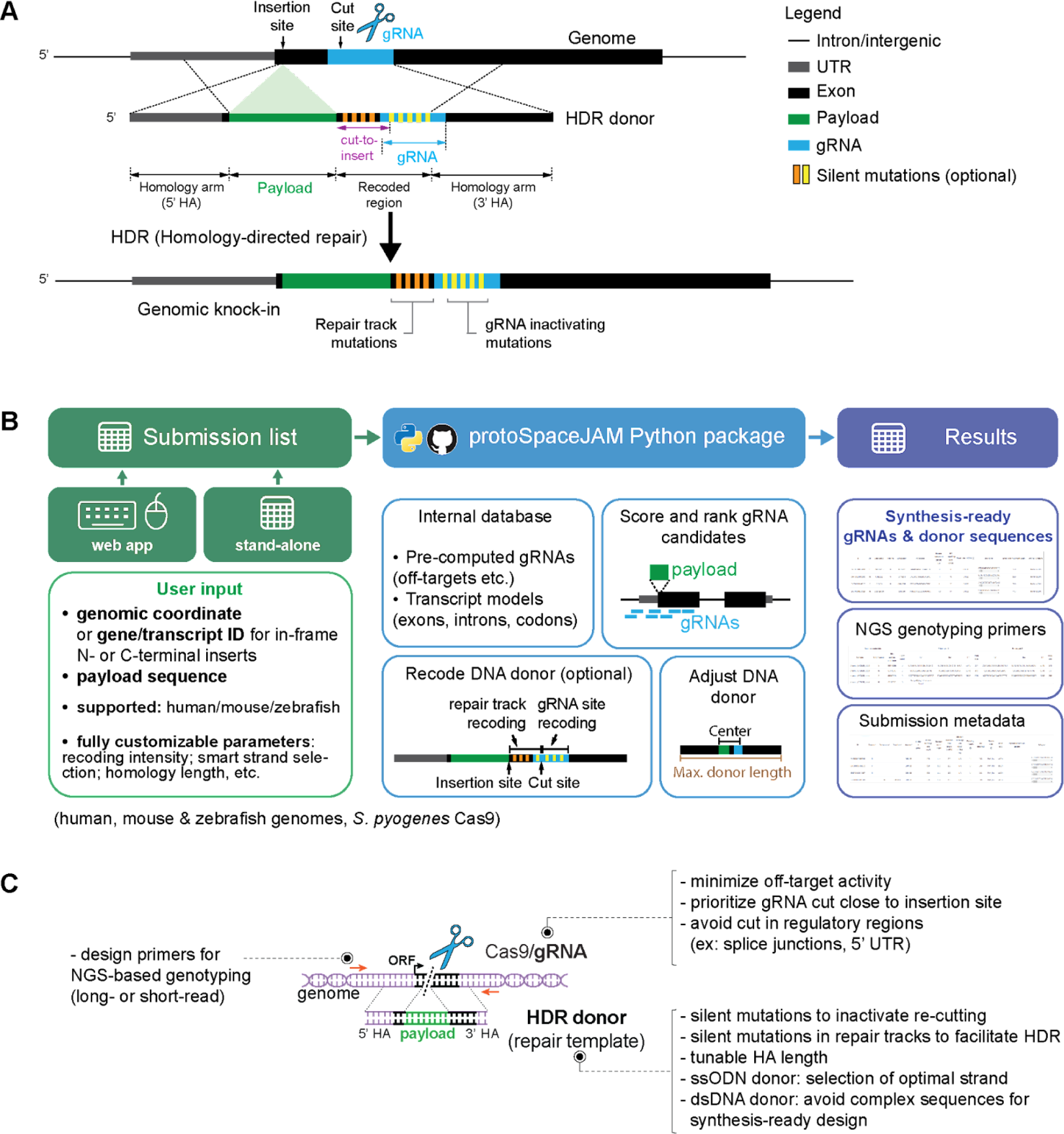
Key concepts, flowchart, and tunable parameters in protoSpaceJAM. **(A)** Key concepts for CRISPR knock-in design. For each insertion, a gRNA (blue) targeting a genomic region and an HDR donor sequence that templates payload (green) integration must be designed. To increase knock-in efficiency, the HDR donor may contain silent mutations to protect the HDR product from further CRISPR-induced DNA strand breaks (“re-cutting” mutations, yellow stripes) and silent mutations in the “cut-to-insert” region between the double-strand break and payload integration sites (“repair track” mutations, orange stripes). Recoded regions are not considered part of the effective homology arms. **(B)** Flowchart of protoSpaceJAM components. protoSpaceJAM can be used either as a standalone Python package or via a user-friendly web app. **(C)** Tunable design parameters in protoSpaceJAM.

The design of a CRISPR/Cas HDR-based insertional knock-in experiment (subsequently referred to as “knock-in”) is conceptually simple and involves two key components. For each edit, a guide RNA sequence (“protospacer”) targeting the desired genomic region must be chosen, and the sequence of a HDR donor template must be constructed (Figure 1A). In recent years, a set of design rules for both gRNA and donor has been established to optimize knock-in efficiency (13–18). This creates an opportunity to develop an algorithm to programmatically design reagents and streamline knock-in experiments. Because of the large breadth of applications of knock-in across genomic, cell biology, or clinical research, such a design algorithm would benefit a large community. Ideally, this algorithm would be 1) accessible and simple to use, enabling researchers from various fields to harness and utilize knock-in technologies; 2) customizable, giving the user full control over the parameters used in the design choices and tailored for each application; and 3) fully open-source, with well-documented source code built in a modular format so that parts of the algorithm could be easily re-used or modified by others to pave the way for new applications.

Here, we present protoSpaceJAM, an easy-to-use, customizable, open-source, and scalable algorithm for CRISPR/Cas9 insertional knock-in. protoSpaceJAM utilizes a state-of-the-art set of rules for gRNA and HDR donor design and enables the user to fine-tune parameters for each design (including the length of homology arms, the introduction of recoding mutations such as silent mutations and mutations in non-coding sequences, or the avoidance of specific sequence features), while providing sensible default options for novice users. protoSpaceJAM allows users to choose between double-stranded DNA (dsDNA) and single-stranded oligodeoxynucleotide (ssODN) forms of HDR donors, with rules tailored to each donor type to optimize knock-in efficiency and/or facilitate chemical DNA synthesis. A companion algorithm, GenoPrimer, designs sequencing primers to simplify the genotyping of edited alleles by next-generation sequencing. protoSpaceJAM is currently built for experiments utilizing the widely used *S. pyogenes* Cas9 nuclease with an NGG Protospacer Adjacent Motif (PAM), and the tool supports insertional knock-in applications in which a payload sequence can be inserted into any coordinate in the human, mouse, or zebrafish genomes (as opposed to mutational knock-in applications where a native sequence is replaced by another). Documented source code is available (github.com/czbiohub-sf/protospacejam) to enable and facilitate the introduction of other desired features. Finally, protoSpaceJAM is available as a stand-alone Python package as well as a user-friendly web platform (protospacejam.czbiohub.org) to catalyze broader accessibility and usage.

## MATERIAL AND METHODS

### Genome datasets

The following reference genomes were used: GRCh38 for human, GRCm39 for mouse, and GRCz11 for zebrafish. Genome annotations for human, mouse, and zebrafish were downloaded as GFF3 files from ftp.ensembl.org/pub/release-107. Custom scripts (github.com/czbiohub-sf/protoSpaceJAM/blob/main/protoSpaceJAM/precompute/scripts/extract_gene_models_info.py) were used to parse the GFF3 files to extract and store the boundaries of UTRs, exons, and introns as genomic coordinates for all annotated transcripts. The reading frame of each position in coding sequences was extracted and stored for every annotated transcript. Sequences near splice junctions were extracted using scripts available at github.com/duopeng/JuncSeq and sequence logo plots were generated using weblogo v3.7.12 (19).

### gRNA search and off-target score calculation

protoSpaceJAM uses precomputed gRNA information to reduce processing time. Precomputed gRNAs for the human, mouse, and zebrafish genomes are available at github.com/czbiohub-sf/protoSpaceJAM#download-and-unzip-pre-computed-data. The computer code and workflow to precompute gRNA for any given genome is available at: github.com/czbiohub-sf/protoSpaceJAM/tree/main/protoSpaceJAM/precompute. Briefly, all possible *S. pyogenes* Cas9 protospacers (NGG PAM) are enumerated for the entire genome. Code from the CRISPOR tool (20) was adapted to calculate target specificity score for each protospacer as follows. First, each protospacer sequence is aligned to the genome using the Burrows-Wheeler Aligner (bwa) (21), allowing up to four mismatches. The non-default parameters employed in this alignment process included “-o 0”, “-m 2000000”, “-n 4”, “-k 4”, “-N”, and “-l 20”. Second, genomic off-target matches are analyzed for the presence of PAMs immediately to the 3’ of the protospacer considering the PAMs NGG, NAG, and NGA. Third, for each off-target match including a PAM, the MIT score (13) is computed. Finally, an aggregated MIT score is computed for each protospacer using the following formula: 100 / (100 + sum of off-target MIT scores).

### gRNA scoring

The strategy for gRNA scoring is described in detail in the main text. All corresponding code can be found at github.com/czbiohub-sf/protoSpaceJAM/blob/main/protoSpaceJAM/util/utils.py.

### Primer design with GenoPrimer

Amplification primers for genotype analysis were designed using GenoPrimer, a custom-built open-source pipeline available at github.com/czbiohub-sf/GenoPrimer. Briefly, the pipeline takes a genomic coordinate position as input (typically the CRISPR knock-in edit site), and it then extracts 430 or 4000 base pairs (bp) of genomic sequences centered on the input coordinate for short- or long-read mode, respectively. From the extracted genomic sequence, candidate primers are enumerated and filtered with Primer3 (22) using a set of thermodynamics filters, a positional filter, and a product size filter. The thermodynamics filters are default for Primer3 and include but are not limited to the following: minimum, optimum, and maximum melting temperatures (Tm) of 57°C, 60°C, and 63°C, respectively; a maximum difference of 3°C in Tm; GC content between 20% and 80%; and primer lengths of at least 18 bp, with an optimum of 20 bp and a maximum of 25 bp. Primer-dimers are minimized by application of default thresholds for both 3’ self-complementary and 3’ pair-complementary binding. The positional filter was configured using the “SEQUENCE_EXCLUDED_REGION” directive to remove primers too proximal to the CRISPR knock-in edit site, with the minimum distance set to 100 bp and 1000 bp for short- and long-read modes, respectively. The PCR product size filter is configured using the “PRIMER_PRODUCT_SIZE_RANGE” directive, and it is set to 250–350 bp and 3300–3700 bp for short- and long-read modes, respectively.

Candidate primers that successfully pass all three filters imposed by Primer3 are further analyzed for unintended PCR products in the target genome. All possible annealing sites in the genome are identified using Bowtie (23) by default, with the following custom parameters: “-k 1000” and “-v 3”. Whereas Bowtie limits alignments to a maximum of three mismatches, BLAST is an alternative mapping program that allows more than three mismatches at the cost of computational speed, this option can be selected by passing the argument “--aligner BLAST” to GenoPrimer with the following custom parameters: “-task blastn-short”, “-max_hsps 2000”, “perc_identity 75”. Annealing sites are removed if the 3’ end of the primer sequence does not match the genome. Unintended PCR products are identified when a primer pair possesses annealing sites on opposite DNA strands with their 3’ ends facing each other, with a maximum predicted amplicon size set to 6 kb by default. GenoPrimer checks whether unintended PCR products can be formed between forward + reverse, forward + forward, and reverse + reverse primers of the same pair; if so, the primer pair is removed from the list of candidates. If no successful primer pairs can be found, a series of six attempts will be made to find the next-best primer pairs. In the first, second, and third attempts, the maximum difference in Tm between paired primers is increased by 1°C. In the second, fourth, and sixth attempts, the upper limit of the PCR product size is increased by 80 bp and 300 bp for short- and long-read mode, respectively. To accommodate the increased PCR product size range, the extracted genomic sequences are extended on both ends by 40 bp and 150 bp for short- and long-read mode, respectively.

### protoSpaceJAM Python package and web portal

protoSpaceJAM is available as a pip-installable Python package available at github.com/czbiohub-sf/protoSpaceJAM. Users have the flexibility to fine-tune the underlying algorithm and customize parameters. Specific documentation to do so is available at github.com/czbiohub-sf/protoSpaceJAM/wiki. For example, one can adjust the mathematical formulas responsible for calculating gRNA scoring weights and redefine how synonymous codons are chosen for silent mutations.

Local versions of protoSpaceJAM require precomputed gRNA information, which can be automatically downloaded and set up for the human, mouse, and zebrafish genomes following the installation instructions at github.com/czbiohub-sf/protoSpaceJAM/blob/main/README.md. Users can precompute gRNA information for genomes of their choice following a set of instructions and scripts available at github.com/czbiohub-sf/protoSpaceJAM/tree/main/protoSpaceJAM/precompute.

An interactive web tool is available at protospacejam.czbiohub.org. All source code for the web tool is available at github.com/czbiohub-sf/protoSpaceJAM-portal, which also includes instructions to set up local versions of the interactive web tool for specific applications.

### Cell culture

Human HEK-293T cells (ATCC #CRL-3216) were cultured in Gibco DMEM, High Glucose, GlutaMAX Supplement media (Thermo Fisher scientific, #10566024) with 10% fetal bovine serum (Omega Scientific, #FB-11). The cells were maintained at 37°C and 5% CO_2_ and were passaged upon reaching ∼80% confluency using 0.05% Trypsin-EDTA (Fisher Scientific, #25300120).

### Genome editing with fluorescent payloads and flow cytometry (cf. Figures 2C, 3C, 4D)

For monitoring of HDR efficiency, we inserted sequences encoding fluorescent proteins in HEK-293T cells using methods described in (1). Briefly, ribonucleoprotein (RNP) *S. pyogenes* Cas9/gRNA complexes were prepared in vitro, mixed with HDR donor templates, and electroporated into HEK-293T cells by nucleofection using SF solution (Lonza #V4SC-2096) and the CM-130 program. Five days post-nucleofection, the distribution of fluorescence signal in each target cell population was analyzed by analytical flow cytometry on a FACSymphony instrument (BD Biosciences). Flow cytometry data analysis was performed using the FlowJo software (BD Biosciences). All gRNA and HDR donor sequences used in this study are available in Supplementary Table 1, together with the corresponding numbers of % fluorescent cells.

**Figure 2.**
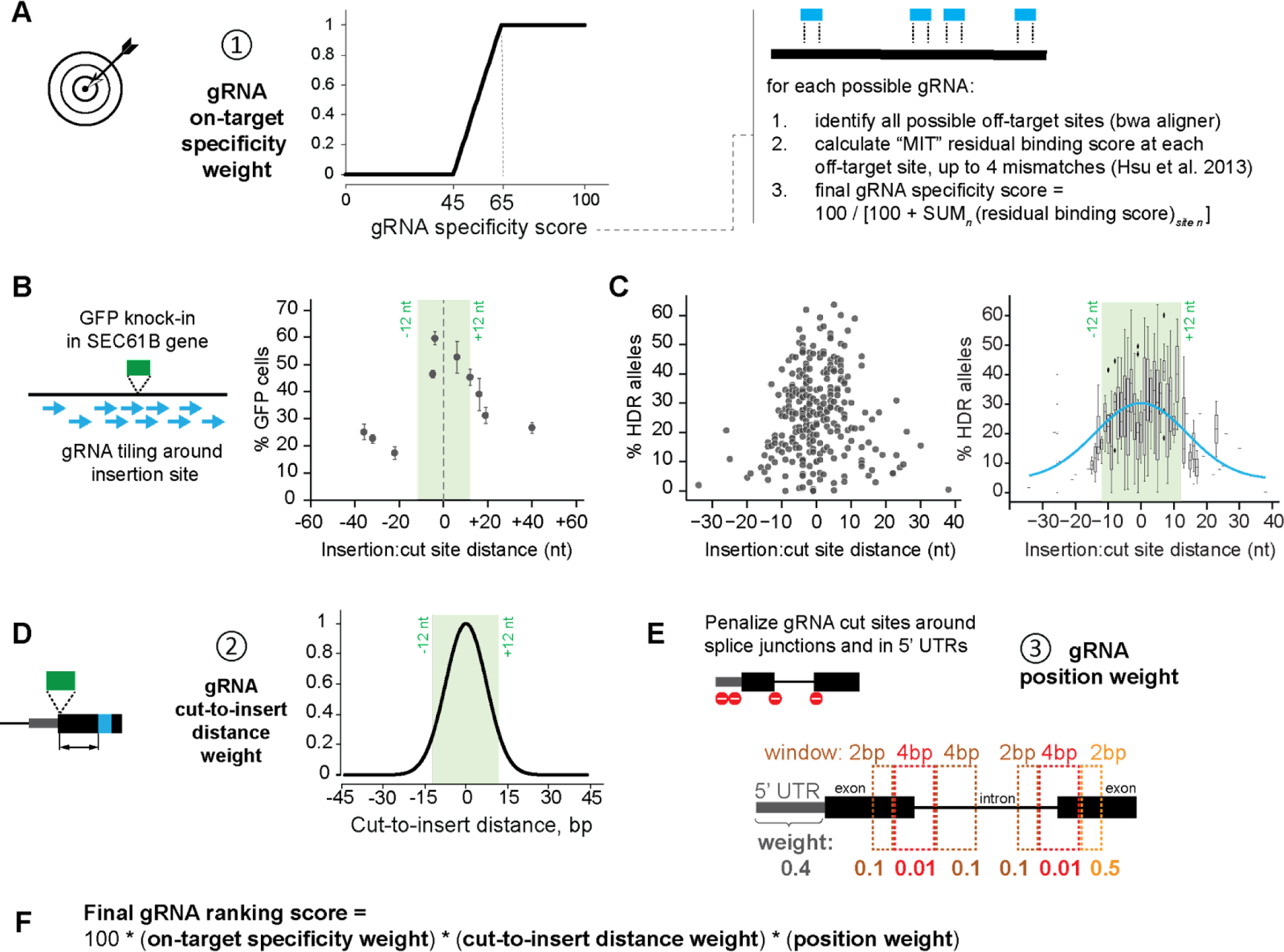
Guide RNA scoring and ranking strategy. **(A)** gRNA specificity is captured by using an “on-target specificity” weight ranging from 0 to 1, where 0 is for specificity scores below 45 and 1 is for specificity scores above 65, and a linear interpolation is made for scores between 45 and 65. Specificity scores are calculated using a variation of the MIT specificity formula described in CRISPOR (30) that considers any possible off-target gRNA binding sites containing up to 4 mismatches to the original protospacer. **(B)** Payload integration efficiency as measured by the integration of a split-GFP payload (2xGFP11) in the SEC61B locus in HEK293T cells using ssDNA donor templates. The percentage of GFP-positive cells is measured by flow cytometry and serves as a read-out of knock-in efficiency. Graph shows average values (filled circles) from n=4 replicates (error bars show standard deviation). **(C)** HDR efficiency as measured by sequence analysis of targeted alleles. The insertion of a split-mNeonGreen payload was characterized across 271 gene loci in HEK293T cells. The rate of HDR for each gene was characterized by deep sequencing of the targeted alleles. The data are presented as a scatter plot (left) or a box plot (right; boxes represent 25th, 50th, and 75th percentiles, and whiskers represent 1.5 times the interquartile range; outliers are shown as diamonds). An unconstrained Gaussian fit to this distribution is presented as a blue line (y = 4.3 + 26.6 × exp[-(x - 2.9)2/360). **(D)** protoSpaceJAM seeks to prioritize gRNAs proximal to the insertion site using a simple “cut-to-insert distance” Gaussian weight between 0 and 1, with a width of around 12 bp (weight = exp[-distance^2/110]). **(E)** Optional “gRNA position” weight between 0 and 1 used by protoSpaceJAM to penalize gRNAs targeting 5’ UTRs or splice junctions. **(F)** protoSpaceJAM ranks all possible gRNAs for each knock-in design using the product of all three “on-target specificity”, “cut-to-insert distance”, and “position” weights in a final composite score.

### Genotype analysis of OpenCell lines (cf. Figure 2C)

The genotype of 271 cell lines from the OpenCell project was analyzed by next-generation sequencing as described in (1). In these experiments, a split-mNeongreen fluorescent payload was inserted in 271 different protein-coding genes in HEK-293T cells by nucleofection of Cas9/gRNA RNPs and single-stranded DNA donors. For each targeted gene, the genotype of a polyclonal pool of ∼20,000 cells was characterized 5 days post-nucleofection in the absence of any selection, so that HDR rates could be directly measured. For each cell pool, genomic DNA was first extracted by cell lysis using QuickExtract DNA Extraction Solution (Lucigen), from which target gene-specific amplicon libraries were prepared using a two-step PCR protocol as described in (1): the first PCR amplifies the target genomic locus and adds universal amplification handle sequences, while the second PCR introduces indexed Illumina barcodes using these universal handles. Barcoded amplicons were analyzed using capillary electrophoresis (Fragment Analyzer, Agilent #DNF-474-0500), pooled, and purified using solid-phase reversible immobilization magnetic beads. Sequencing was performed on an Illumina Miseq V3 platform (2×300 bp paired-end reads) using standard P5/P7 primers. Genotype analysis was performed using CRISPRESSO2 (24) to quantify HDR rate for each target gene (defined as the percentage of HDR alleles out of all non-wild type alleles sequenced, to normalize for the cutting efficiency of each gRNA). gRNA, HDR donor, and primer sequences and genotype analysis for all targets are found in Supplementary Table 2.

### GenoPrimer amplification test (cf. Figure 5B)

To test GenoPrimer’s ability to design amplification primers for genotype analysis, we designed primer pairs for 94 separate genes in short-read mode. Products were amplified from wild-type genomic DNA purified from wild-type HEK-293T cells (New England Biolabs #T3010S) using a touch-down PCR strategy. 40-µL PCR reactions were set using 2x KAPA HiFi Hotstart reagents (Roche #KK2602) with 10 ng genomic DNA, 2 µM of each primer, and betaine to 1M final concentration. PCR conditions: 95°C for 3min; 2 cycles of {98°C for 20s, 72°C for 15s, 72°C for 20s}, 2 cycles of {98°C for 20s, 71°C for 15s, 72°C for 20s}, 2 cycles of {98°C for 20s, 70°C for 15s, 72°C for 20s}; 2 cycles of {98°C for 20s, 69°C for 15s, 72°C for 20s}; 2 cycles of {98°C for 20s, 68°C for 15s, 72°C for 20s}; 2 cycles of {98°C for 20s, 67°C for 15s, 72°C for 20s}; 2 cycles of {98°C for 20s, 70°C for 15s, 66°C for 20s}; 2 cycles of {98°C for 20s, 65°C for 15s, 72°C for 20s}; 2 cycles of {98°C for 20s, 70°C for 15s, 64°C for 20s}; 2 cycles of {98°C for 20s, 63°C for 15s, 72°C for 20s}; 10 cycles of {98°C for 20s, 62°C for 15s, 72°C for 20s}; then 72°C for 1 min (final extension); 4°C final. After PCR, the size distribution of amplicons for each target gene was characterized by quantitative capillary electrophoresis (Fragment Analyzer, Agilent #DNF-474-0500). All primer sequences for the GenoPrimer test are found in Supplementary Table 3.

## RESULTS

### Anatomy of knock-in design and overview of protoSpaceJAM

protoSpaceJAM is a design tool currently developed for insertional HDR-based knock-in applications with *S. pyogenes* Cas9 nuclease. In these applications, the genomic insertion of a functional payload is templated by a HDR donor containing the payload flanked by sequences called “homology arms” that are homologous to the desired insertion site (Figure 1A). For each insertion, two separate components must be designed: a gRNA targeting the genomic region of interest, and the HDR donor sequence itself. In cases where the original protospacer might be preserved within one of the homology arms, introduction of silent “recoding” mutations may be desirable to inactivate gRNA binding and re-cutting of the knock-in allele (25) and/or to re-code the genomic portion located between the desired insertion site and the Cas9/gRNA cut site (26) (see Figure 1A; the rationale for recoding mutations is further explained in the next sections).

protoSpaceJAM employs a simple input: the user specifies a genomic insertion site and a payload sequence (Figure 1B). Currently, designs for human, mouse, and zebrafish genomes are supported, and design speed is maximized by building an internal database of pre-computed off-target information of all possible *S. pyogenes* Cas9 protospacers (NGG PAM) in each genome. Because one of the main applications of knock-in is to insert functional tags in-frame of a protein-coding gene (e.g., for fluorescent protein tagging), the user can also specify a transcript ID for a gene of interest and an N- or C-terminal insertion point. Explicit genomic coordinates can be used for more flexible designs.

The goal of protoSpaceJAM is to streamline the design of both gRNA and donor sequences using a biologically-informed set of rules that are fully described in subsequent sections (summarized in Figure 1C). Importantly, all parameters are fully controlled by the user to maximize utilization across many different applications. Multiple knock-in designs can be handled in parallel by building a submission list, which contains insertion position and payload information as well as the parameter details to be used for each design (Figure 1B). All documented source code is openly available at github.com/czbiohub-sf/protoSpaceJAM, and the tool can be used either as a pip-installable standalone Python package or via a user-friendly web interface at protospacejam.czbiohub.org.

### gRNA selection rules

The first step in knock-in design is the selection of a desirable gRNA for optimal efficiency and specificity. To rank all candidate gRNAs for optimal selection, protoSpaceJAM uses a composite ranking score that weighs the on-target specificity of each candidate, the distance between cut and insertion sites, and the position of the gRNA with respect to important gene expression regulatory sequences, namely 5’ untranslated regions (UTRs) and splice sites (Figure 2 A-E).

#### Specificity weight

A key feature of each gRNA is its specificity for a single genomic site, as Cas9 might cut at off-target genomic sequences that share significant homology to a given protospacer (27, 28). Multiple scoring algorithms have been developed to predict gRNA specificity (reviewed in (29)), including the widely used scoring strategy originally developed by Hsu et al. and referred to as the MIT score (13). protoSpaceJAM uses a variation of the MIT specificity score described in CRISPOR (30) that considers any possible off-target gRNA binding sites containing up to 4 mismatches to the original protospacer. Off-target modification frequencies drop significantly for gRNAs with specificity scores above 45, and they become minimal for scores above 65 (29). For ranking purposes, protoSpaceJAM uses a simple rule to capture gRNA specificity using an “on-target specificity weight” between 0 and 1, where 0 is assigned to gRNAs with a specificity score below 45 (to severely penalize non-specific guides), 1 is for specificity scores above 65, and a linear interpolation is made for scores between 45 and 65 (Figure 2A).

#### Cut-to-insert distance weight

For knock-in applications, many studies have shown that the distance between the Cas9/gRNA cut site and the desired insertion point is a key parameter that governs integration efficiency (10, 15, 25, 31, 32). The further the genomic cut site is from the insertion site, the greater the probability that DNA repair might resolve without payload insertion (because homology to the genome that exists in the donor region between cut and insert will allow DNA repair to resolve prematurely, Suppl. Figure 1A). To test the relationship between integration efficiency and cut-to-insert distance, we measured the integration of a fluorescent payload in the human SEC61B protein-coding gene while tiling gRNAs across the insertion point. Integration efficiency (measured by the percentage of fluorescent cells detected) dropped sharply as a function of cut-to-insert distance, with distances over 12 bp exhibiting the greatest decrease (Figure 2B). Similar relationships have been reported in the literature (33). To further characterize the relationship between HDR efficiency and cut-to-insert distance, we analyzed data from the OpenCell project (1) in which a short fluorescent payload was inserted in a large number of human protein-coding genes, fortuitously spanning a range of cut-to-insert distances. For 271 insertions (randomly chosen), the targeted allele was sequenced from edited cell pools and the rate of HDR was quantified (see Material and Methods). This analysis also shows a reduction of HDR efficiency as cut-to-insert distance increases (Figure 2C). As a consequence of these results, protospaceJAM uses a scoring weight to prioritize gRNAs with cut sites proximal to the insertion site. By default, protospaceJAM uses a “cut-to-insert distance” Gaussian weight between 0 and 1, with a width centered around 12 bp (Figure 2D).

#### Additional positional weight

Targeting Cas9/gRNA cuts within important regulatory sequences such as 5’ UTRs or splice junctions should be avoided because the subsequent need to introduce recoding mutations around the cut site (see below) might impact these regulatory regions and alter the endogenous expression of the target gene. 5’ UTRs contain ribosome-binding sequences and play a key role in controlling gene expression (34), while splice junctions contain residues that are universally conserved and cannot be modified (Suppl. Figure 2). Therefore, we defined a “gRNA position weight” between 0 and 1 to penalize gRNAs targeting these regions (see Figure 2E). Because this optimization strategy is only relevant to some applications, using the positional weight is optional.

#### Final weight calculation and gRNA scoring

For each knock-in design, protoSpaceJAM ranks all possible gRNAs using the product of the “on-target specificity”, “cut-to-insert distance”, and “position” weights in a final composite score (Figure 2F). The user can specify the desired number of gRNAs to be returned for each design, prioritized according to their composite score. The detailed weight information for each guide is specified in protoSpaceJAM’s table output. Overall, the goal of protoSpaceJAM is to nominate gRNAs that are specific and likely to achieve high knock-in rates while minimally affecting the genetic regulatory patterns of endogenous genes. In our open-source code, the parameters that govern gRNA selection are explicitly annotated and organized as separate modules. Users who would like to customize or add selection criteria can easily modify the corresponding modules.

Note that beyond positional prioritization, protoSpaceJAM does not score the predicted editing activity of gRNAs. Although gRNA activity scores have been developed (29), these have been shown to be application-dependent and to often correlate with the expression levels of gRNA sequences under specific promoter conditions (e.g., using U6 or T7 promoters; see (29)). Because these scores do not reflect gRNA activity under universal conditions, and because gRNA expression is not a limiting factor in common protocols that use direct delivery of purified CRISPR/Cas complexes (35, 36), we currently do not use activity prediction in our calculations.

### Mutational recoding in the HDR donor

protoSpaceJAM supports the optional introduction of silent “recoding” mutations in two key regions of the HDR donor (Figure 1A). The first region is the Cas9/gRNA binding site, which may still be present in the homology arm sequences when payload insertion does not destroy the original protospacer. In such cases, knock-in could be impaired if Cas9 either cuts the donor itself during the delivery of reagents in the cell or re-cuts the knock-in allele after DNA repair (25, 37, 38). This would respectively decrease donor availability or introduce unwanted genomic modifications, negatively impacting knock-in efficiency overall. A well-established practice is therefore to introduce silent mutations to inactivate the gRNA binding site within the HDR donor (25, 37, 38). protoSpaceJAM uses the Cutting Frequency Determination (CFD) scoring framework established by Doench and colleagues to predict the impact of individual protospacer and PAM mutations on the Cas9/gRNA cutting potential (14). For each gRNA, protoSpaceJAM identifies the fewest mutations that would bring the maximal CFD score in the donor sequence below a user-defined threshold (default: 0.03). Because the payload sequence itself may by chance contain sequences homologous to the Cas9/gRNA binding site, all positions within the donor (including payload and payload/genome junctions) are considered in this calculation. When recoding within a protein-coding sequence, only silent mutations are used, leveraging maximal sequence divergence between synonymous codons while excluding rare codons. When recoding within a non-coding region, mutations are introduced in up to one of every three bases (Figure 3A). No recoding is allowed in the immediate vicinity of splice junctions, to maintain universally conserved sequence motifs (Suppl. Figure 2). For maximal flexibility, the user can decide whether recoding should prioritize mutations in the PAM region (default) or within the protospacer itself.

**Figure 3.**
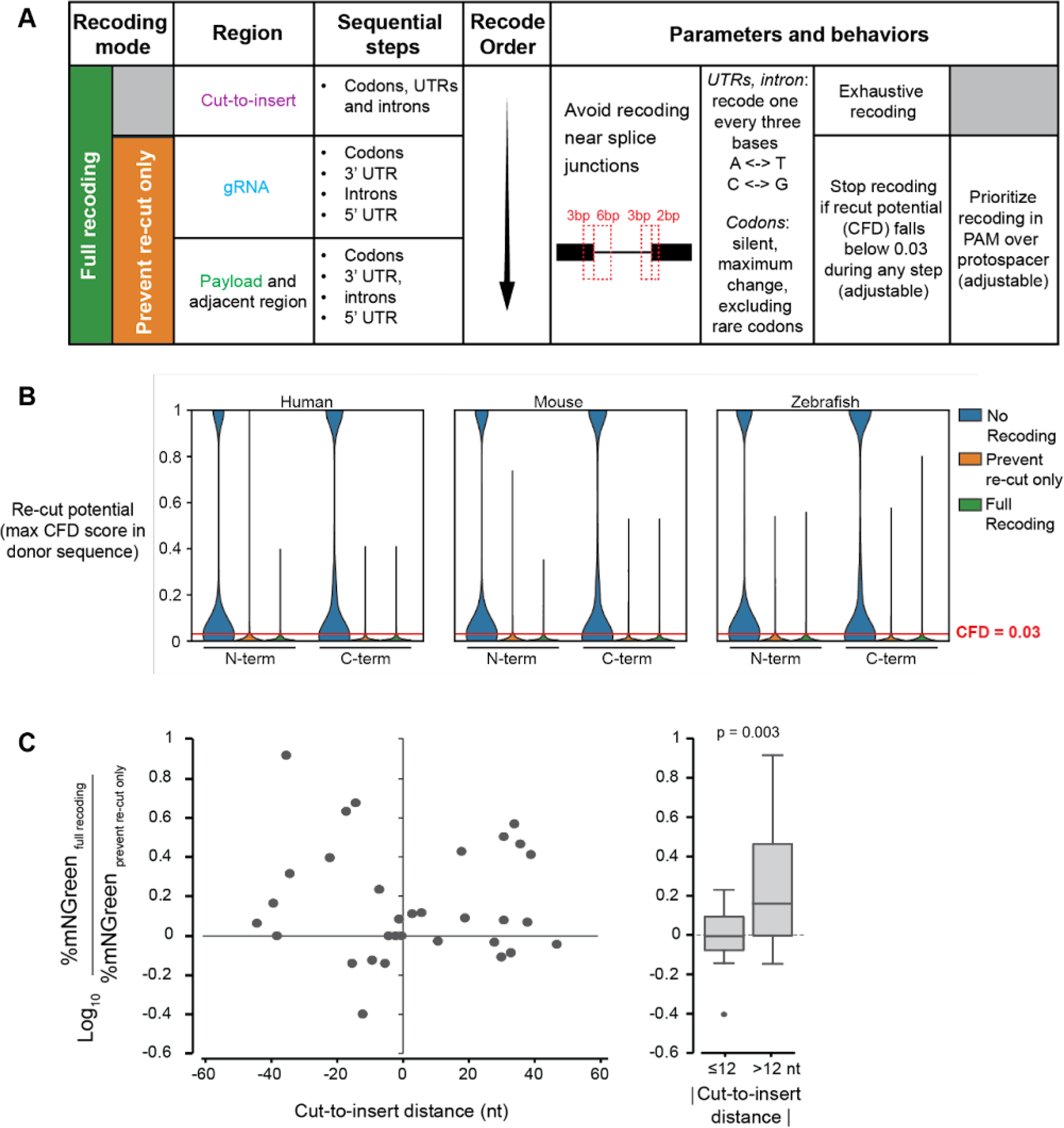
Mutational recoding in the HDR donor. **(A)** Recoding options in protoSpaceJAM. Two recoding levels are available: “prevent re-cut only” (orange) introduces silent mutations around the Cas9/gRNA binding site to prevent re-cutting of the knock-in allele, while “full recoding” (green) additionally introduces silent mutations in the cut-to-insert region to prevent DNA repair from resolving before incorporation of the payload sequence. **(B)** Recoding brings the re-cut potential below the desired threshold (CFD < 0.03) in 97.6%, 97.8%, and 97.2% of knock-in alleles for insertion of a split-mNeonGreen payload (1) at the N- or C-terminus of all canonical protein-coding transcripts in the human, mouse, or zebrafish genomes (n = 19643, 24714, and 25406, respectively). **(C)** “Repair track” recoding in the cut-to-insert region affects knock-in efficiency of a split-mNeonGreen payload across 32 human protein-coding genes for which top-scoring gRNAs were located at various cut-to-insert distances. The percentage of GFP-positive cells is measured by flow cytometry and serves as a read-out of knock-in efficiency.

Recoding can be implemented separately to increase the efficiency of knock-in when having to perform Cas9/gRNA cuts at a distance from the insertion site. In such cases, introducing silent mutations in the cut-to-insert region prevents the DNA repair tracks from resolving repair before reaching the payload sequence (Suppl. Figure 1A), thereby increasing the rate of payload insertion (16, 26). protoSpaceJAM supports recoding within the cut-to-insert region, following the rules outlined above for coding and non-coding sequences and excluding recoding at splice junctions (Figure 3A).

Importantly, the level of recoding for each design is controlled by the user and can be tuned to one of three behaviors: 1) “no recoding”, where no mutations are introduced besides the insertion of the payload itself; 2) “prevent re-cut only”, where mutations are only introduced to inactivate gRNA binding anywhere in the donor sequence; and 3) “full recoding”, where mutations are introduced to inactivate gRNA binding in the donor and repair track mutations are introduced in the cut-to-insert region. By default, we recommend using full recoding. The advantage of recoding is illustrated in Figures 3B-3C. Figure 3B shows the potential for Cas9/gRNA re-cutting of knock-in alleles for GFP insertion at the N- or C-terminus of all protein-coding genes in the human, mouse, or zebrafish genomes, as predicted by the maximal residual CFD score within each donor sequence. In the absence of recoding, 53.6%, 55.4%, and 61.0% of knock-in sequences exhibit significant re-cutting potential in human, mouse, and zebrafish, respectively (Figure 3B, “no recoding”; note that when gRNAs proximal to the insertion sites can be chosen, the insertion of the payload by itself may be enough to impact the protospacer and prevent re-cutting). The introduction of silent recoding mutations brings the re-cutting potential below the desired threshold in 97.6%, 97.8%, and 97.2% of the sequences in human, mouse, and zebrafish, respectively (Figure 3B). In addition, adding repair track mutations in the cut-to-insert region increases knock-in efficiency for gRNAs with large cut-to-insert distances (Figure 3C). To verify this, we measured the integration efficiency of a fluorescent mNeonGreen payload across 32 human protein-coding genes for which top-scoring gRNAs were located at different cut-to-insert distances, comparing designs using the “full recoding” vs “prevent re-cut only” modes (Figure 3C, left). The full recoding option led to significantly higher integration efficiency (as measured by the percentage of fluorescent cells detected) for designs in which the cut-to-insert distance was greater than 12 nt (Figure 3C, right panel; note that gene-specific sequence features beyond the cut-to-insert distance are likely to influence knock-in efficiency). These results mirror previous reports in the literature describing the advantages of recoding the cut-to-insert region to increase knock-in efficiency (26, 38).

### Additional design parameters for dsDNA vs ssODN donors

Double-stranded DNA (dsDNA) and single-stranded oligodeoxynucleotides (ssODNs) are the two main formats for HDR donors (8). ssODNs are widely available through commercial DNA synthesis, are non-toxic, and can be delivered to cells in large amounts to maximize knock-in efficiency (39). However, they are limited to small insertions because their overall length is constrained by the coupling efficiency of chemical DNA synthesis (typically ≤200 nt). dsDNA donors, which can be delivered to cells in the form of DNA plasmids or linear PCR-amplified fragments, are a more universal form of donor that can be used for insertions of any size. However, they are comparatively more toxic (12) and may require molecular cloning for generating the amounts needed for efficient delivery. Increasingly, sequence-verified dsDNA constructs are becoming commercially available at attractive price points, but commercial dsDNA products are typically limited to “simple-to-synthesize” sequences with balanced GC content and devoid of homopolymeric stretches. Because homology arms often contain parts of the non-coding genome, HDR donor sequences might often fall beyond the current limitations for commercial dsDNA synthesis.

A key goal of protoSpaceJAM is to provide the user with “synthesis-ready” donor sequences to streamline the knock-in experimental process. Therefore, the user can choose between two separate donor design modes – dsDNA and ssODN – that use separate design constraints. In dsDNA mode, sequence motifs that might be incompatible with commercial dsDNA synthesis are flagged within the final output table. These flags include homopolymeric runs of 10 or more As and Ts or 6 or more Gs and Cs, and extreme GC content (>65% or <25% GC content globally or >52% difference in GC content between any given 50 bp stretches). We analyzed the presence of these flags in homology arms of 500 bp flanking the N- and C-terminus tagging sites in all canonical transcripts in human, mouse, and zebrafish genomes (39637, 43719, and 55559 sites, respectively). Our results indicated that homopolymeric runs are the leading impediment to successful synthesis of the dsDNA donor (Figure 4A). Therefore, we incorporated a feature to directly trim the length of homology arms to remove homopolymers from the dsDNA donor (Figure 4B). The user controls whether trimming should be performed, as well as the minimum length of homology arms to be retained. To assist users in making informed decisions regarding the minimum length of homology arms, we analyzed 39,637 N- and C-terminus knock-in sites in the human genome. Our findings indicate that DNA donor arms with lengths of 100 bp, 200 bp, and 500 bp have a 94%, 87%, and 70% probability of being free from homopolymers, respectively (Figure 4B). Our trimming algorithm halts once it either meets the minimum length requirement or the sequence becomes free of homopolymers, whichever condition is satisfied first. Trimming can also be used to avoid other user-defined sequence motifs, such as restriction sites for enzymes used in subsequent applications (Figure 4C).

**Figure 4.**
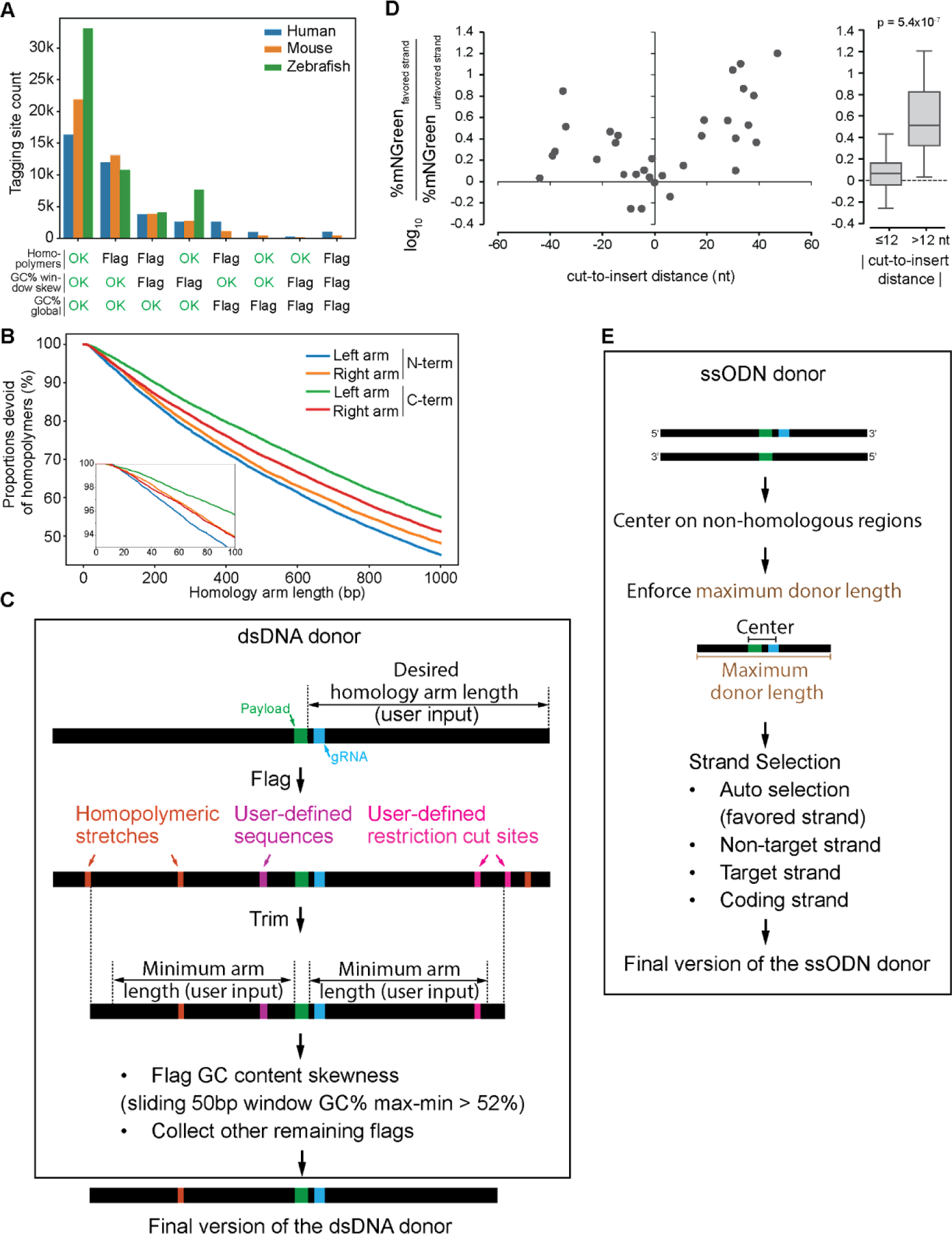
Design parameters for dsDNA and ssODN donors. **(A)** Analysis of 500-bp homology arms flanking the N- and C-terminus tagging sites in all canonical transcripts in human, mouse, and zebrafish genomes indicates that homopolymeric runs are the leading problem that would prevent successful synthesis of the dsDNA donor. Homopolymers are defined by homopolymeric runs of 10 or more As and Ts or 6 or more Gs and Cs; GC% window skew is defined by GC content skewness between any sliding window of 50 bp exceeding 52%; GC% global is flagged when the global GC content is below 25% or above 65%. **(B)** Proportions of homology arms devoid of homopolymers as a function of length. N- and C-terminus insertion sites (n = 39286) in all canonical transcripts in the human genome were analyzed. **(C)** Workflow to remove or flag hard-to-synthesize motifs in dsDNA donor. **(D)** Integration efficiencies of a split-mNeonGreen fluorescent payload across 32 human protein-coding genes for which top-scoring gRNAs were located at various cut-to-insert distances, comparing both strand orientations for each ssODN donor. The percentage of mNeonGreen-positive cells is measured by flow cytometry and serves as a read-out of knock-in efficiency. **(E)** Workflow to design synthesis-ready ssODN donor sequences.

For ssODN synthesis, there is typically no restriction in terms of sequence motifs, but rather in overall length. Therefore, the total length of donors in ssODN mode is capped at a user-defined maximum (default: 200 nt). However, ssODN donors require a choice of polarity for the ssDNA strand to be used. The polarity of the ssODN strand is especially important when using gRNAs with a large cut-to-insert distance, and the most efficient strand orientation depends on whether the cut site is on the 3’ or 5’ side of the insertion site (see Suppl. Figure 1B). This is because ssODN donors predominantly template DNA repair via a synthesis-dependent strand annealing mechanism (26). In this mechanism, 5’-end resection following the Cas9-induced double-strand break exposes single-stranded genomic overhangs that can anneal to complementary sequences in the ssODNs donors, which are subsequently extended by DNA synthesis in the 5’ to 3’ direction (Suppl. Figure 1B). This mechanism induces a polarity in the repair process that strongly favors a specific DNA strand for distal payload integration (Suppl. Figure 1B). By default, protoSpaceJAM automatically selects the polarity of the ssODN strand to be in the favored orientation. To validate this design choice, we measured the integration efficiency of a split-mNeonGreen fluorescent payload across 32 human protein-coding genes for which top-scoring gRNAs were located at different cut-to-insert distances, comparing both strand orientations for each ssODN donor (Figure 4D). The favored strand orientation (as defined in Suppl. Figure 1) led to increased integration efficiency, as measured by the percentage of fluorescent cells, particularly for designs in which the cut-to-insert distance was greater than 12 nt (Figure 4D, right panel). Similar strand preference requirements have been previously described and further support our default design choice (15, 26, 40, 37, 38). To give the user even finer control over the ssODN strand to be used, four other strand selection modes are also available: Cas9/gRNA target vs non-target strand or transcribed vs non-transcribed strand for protein-coding genes and lncRNAs (Figure 4E).

### Genotyping primer design for characterization by next-generation sequencing

After knock-in, the genomic sequence of the edited alleles must be characterized to validate successful editing and the absence of undesired errors introduced during DNA repair. This is often done using next-generation sequencing (NGS) of amplicons covering the edit site and homology arms (41-43). Depending on the sizes of the knock-in insertion and the DNA donor used, genotyping can be performed using short- or long-read sequencing. Amplification must be driven by primers that bind the genome outside of the homology arms used in the knock-in design (Figure 5), so that amplification cannot be templated from the HDR donor itself but must instead represent bona-fide edited alleles.

**Figure 5.**
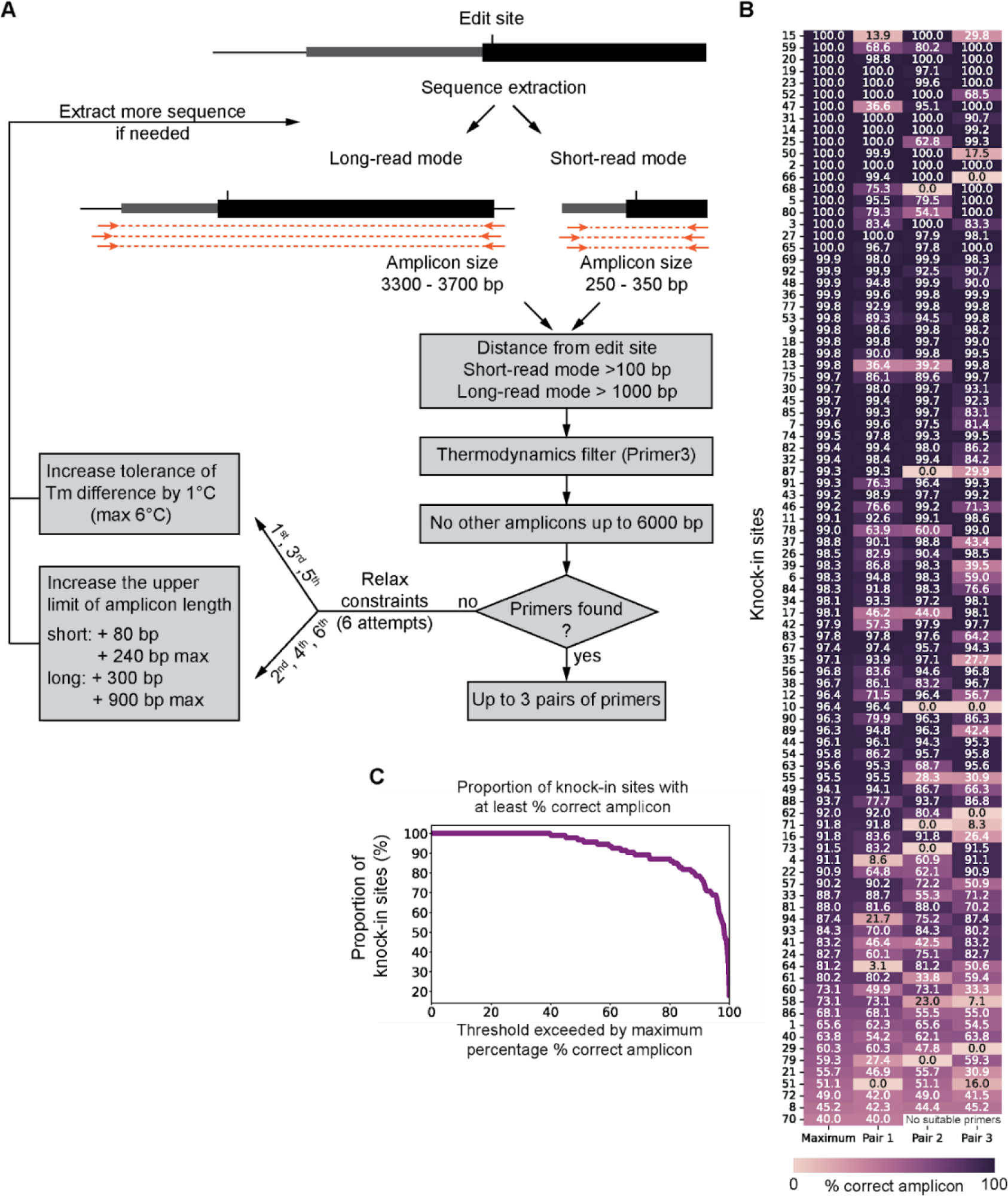
Computational pipeline for design of genotyping primers and experimental validation. **(A)** GenoPrimer predicts primers suitable for amplifying PCR products for short- and long-read sequencing, with amplicon size ranges of 250–350 bp and 3300–3700 bp, respectively. Primer candidates undergo additional screening using thermodynamic filters implemented in Primer3. To check for unintended PCR products, all potential annealing sites in the genome are searched using Bowtie (or optionally BLAST+). Unintended PCR products can be identified when a primer pair has annealing sites on opposite DNA strands with their 3’ ends oriented towards each other. If no candidate primers remain, GenoPrimer employs a strategy of six sequential attempts to find the next-best primer pairs. This is done by gradually increasing the tolerance for differences in melting temperature (Tm) and extending the upper limit of amplicon length. **(B)** Experimental validation. GenoPrimer was used to design short-read amplification primers for 94 separate loci. At least one primer pair matching the criteria outlined in (A) was identified for 93 out of 94 loci (98.9% success rate). Amplification with each primer pair was validated using HEK293T genomic DNA and quantified by capillary electrophoresis. The percentage of amplicons of the correct size over all amplified products is shown for each of the top 3 primer pairs, along with the maximum percentage among all primer pairs for each locus. **(C)** Proportion of knock-in sites for which the percentage of amplicons of the correct size (y-axis) is above a specific threshold (x-axis).

We developed a computational pipeline called GenoPrimer that enables rapid primer design for genotyping edited sites (Figure 5). GenoPrimer is fully integrated into the protoSpaceJAM web tool, allowing users to easily access and utilize this function. Additionally, we provide an open-source standalone version, available at github.com/czbiohub-sf/GenoPrimer. GenoPrimer operates in two distinct modes: short-read and long-read. Short-read mode returns primer for 250-350 bp amplicons, and long-read mode returns primers for 3300–3700 bp amplicons. Candidate primer pairs are selected using three primary filters: 1) a minimum distance from the edited site of 100 bp for short reads or 1000 bp for long reads, to ensure that at least one primer binds outside the homology arms (assuming a combined 5’+ 3’homology length of ≤200 nt for ssODN donors and ≤2000 bp for dsDNA donors); 2) compliance with default thermodynamic criteria used by Primer3 (44) to ensure optimal PCR efficiency (e.g., similar Tm and low probability of binding and formation of primer dimers and secondary structures); and 3) absence of off-target amplicons under 6000 bp, which can arise when both primers in the pair bind non-specifically in opposite orientations on the same off-target chromosome. To identify off-target binding sites, users can select either the Bowtie aligner (23), which is computationally fast but limited to sites with up to 3 mismatches, or BLAST+ (45), which is slower but allows more mismatches. In cases where no desirable candidates can be found, the default design constraints on ΔTm and amplicon length are iteratively relaxed to nominate alternative primer pairs (Figure 5A). When GenoPrimer successfully identifies suitable primers, it outputs up to three pairs of primers along with their PCR product sizes for unedited alleles and annealing temperatures. We used GenoPrimer to compute genotyping primers using short-read mode for N- and C-terminus insertion sites in all canonical open reading frames in the human, mouse, and zebrafish genomes; primers that fit the selection criteria were successfully identified for 38190 of 39200 (97.4%), 41699 of 43579 (95.7%), and 48688 of 50724 (96.0%) sites, respectively. The same computation using long-read mode yielded primers for 98.4%, 97.6%, and 95.0% of sites, respectively. Next, we experimentally tested primers computed for 94 knock-in sites across different human genes (three primer pairs per site). Of these, 93 sites (98.9% of total) yielded primers that fit the selection criteria, with the only exception resulting from an exact sequence duplication between chromosomes that prevents identifying primers devoid of off-target binding sites (CDC26 C-terminus: chr9:113267016-113267441 duplicated on chr7:129409866-129410291). For all 93 sites, at least one primer pair produced an amplicon matching the expected size, as determined by quantitative capillary electrophoresis (Figure 5B). In 96.7% of the cases, the fraction of amplicons of the correct size exceeded 50% of all amplified products, and in 87% of the cases, correct amplicons exceeded 80% of all amplified products (Figure 5C).

### Interactive web tool at protospacejam.czbiohub.org

We developed a user-friendly, interactive, flexible, and fast web-based tool to streamline the use of protoSpaceJAM and GenoPrimer (Figure 6). This tool effectively eliminates the need for command-line prompts, enhancing accessibility for users of all skill levels. It utilizes an intuitive step-by-step navigation system, guiding users through each stage of the process – configuration, verification, execution, and download of CRISPR knock-in designs.

**Figure 6.**
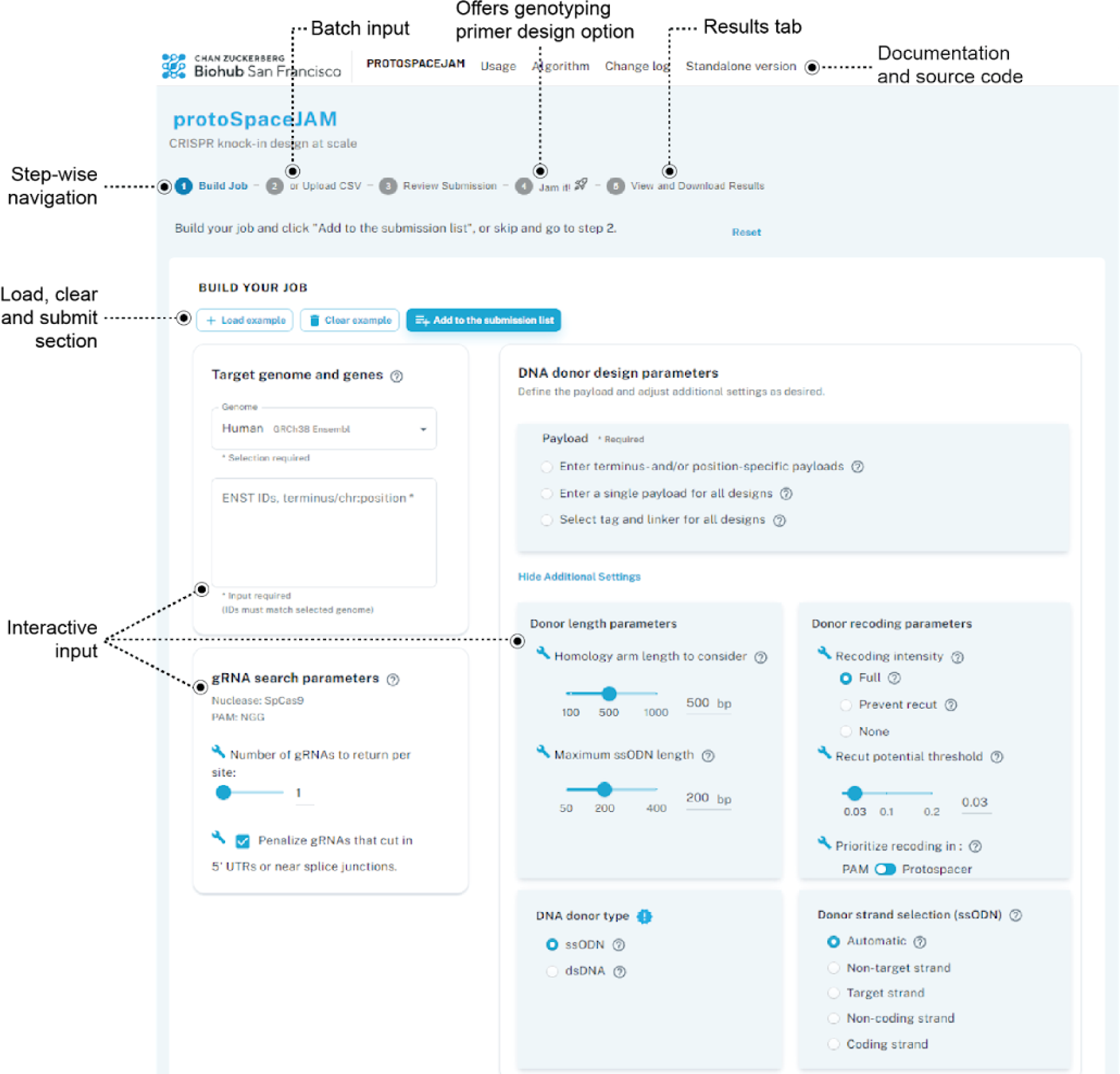
Annotated screenshot from interactive web tool at protospacejam.czbiohub.org.

Knock-in design jobs on protospacejam.czbiohub.org are built around four separate phases (full manual available at czbiohub-sf.github.io/protoSpaceJAM/index.html). In the “*Build your job*” phase, users can interactively parameterize their design requests online. Alternatively, design features can be input from a user-prepared file through the “*Upload csv*” function. Notably, these two steps can continuously feed the same submission list. This enables different designs using different parameters to be processed in the same job. Once a desired submission list has been populated, that list can be reviewed and confirmed in the “*Review submission*” phase. Next, the design job can be launched in the “*Jam it!*” phase, with the option to also design genotyping primers suitable for either short- or long-read sequencing. Finally, in the “*Results*” phase, users can view gRNAs, donors, and primers (if applicable) for each design and download both the results and the submission list, which includes parameterization information, as csv files (Figure 1B).

Users can access documentation, in the style of ReadTheDocs, by following the navigation links situated at the top of the page. For additional assistance, hover tooltips offer immediate contextual explanations of various parameters.

The full source code of our interactive web portal is available at github.com/czbiohub-sf/protoSpaceJAM-portal. Users have the flexibility to set up personalized instances of the web portal, supporting Ensembl-compatible genomes, on their own computing infrastructure.

## DISCUSSION

### Comparison to existing CRISPR/Cas design tools

We developed protoSpaceJAM to fill an existing gap in the rich ecosystem of CRISPR/Cas design tools. While many algorithms have been developed to select gRNAs for diverse applications (reviewed in Figure 7), few support the design of both gRNA and HDR donors for HDR-based knock-in. In addition, the existing tools for knock-in design are all (to our knowledge) developed by commercial DNA synthesis companies and do not have publicly available source code, limiting their re-use by the community (Figure 7). protoSpaceJAM differentiates itself by being the first fully open-source design tool specifically developed for CRISPR/Cas knock-in. In addition, protospaceJAM uses a rich set of explicit and biologically-informed design parameters for gRNA selection and optimal HDR donor construction. In particular, protoSpaceJAM is unique in its ability to 1) penalize gRNAs targeting important regulatory sequences (Figure 2C), 2) allow users to choose between different levels of HDR donor recoding with silent mutations, including recoding in the cut-to-insert region to increase knock-in efficiency (Figure 3C), and 3) use different practical design rules for dsDNA vs. ssODN donor design, including automated selection of optimal strand orientation for ssODN donors and the design of synthesis-ready sequences for dsDNA donors (Figure 4E). In addition, we provide GenoPrimer as a companion algorithm to streamline the design of genotyping primers for both short- and long-read sequencing (Figure 5).

**Figure 7.**
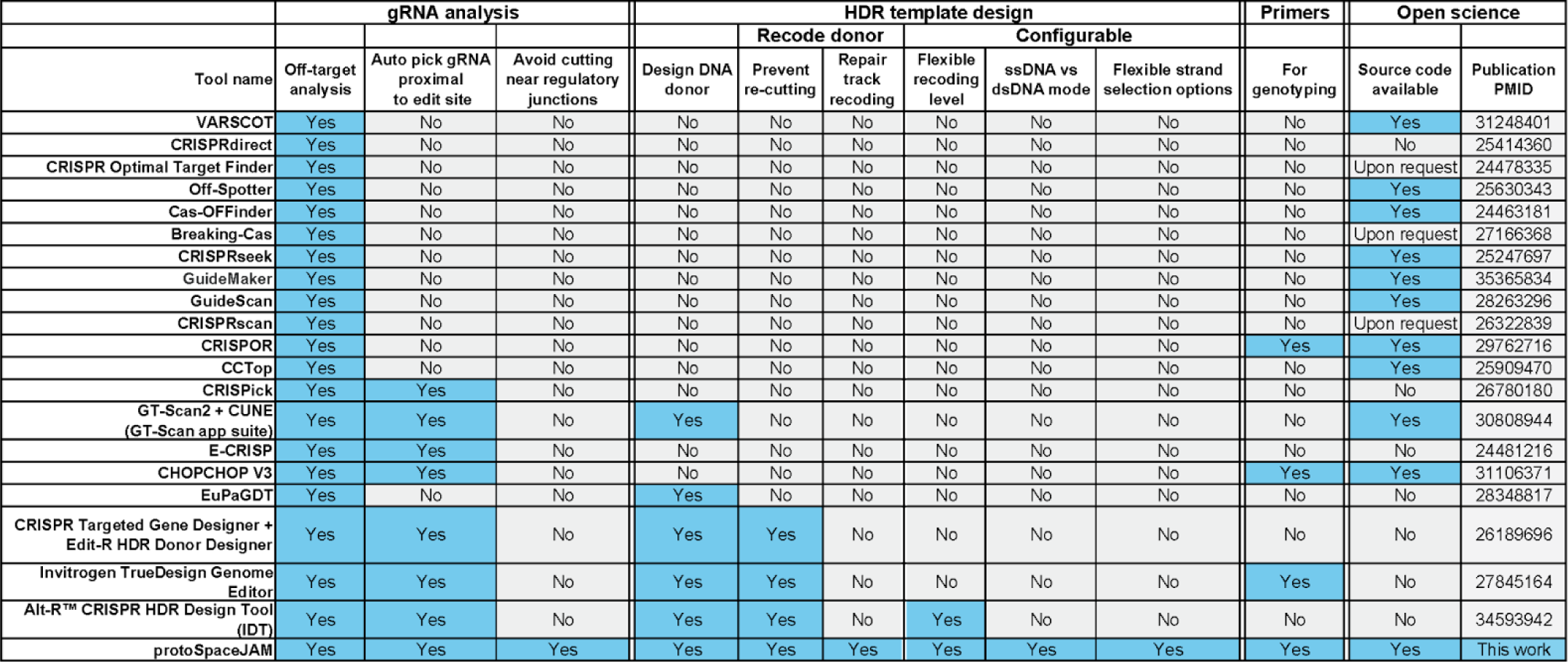
Comparison of protoSpaceJAM with published CRISPR/Cas design tools.

Altogether, we have developed an integrated open-source, modular, and fully customizable design platform to accelerate the design and increase the success rate of HDR-based CRISPR/Cas knock-in experiments. The features included in protoSpaceJAM are informed by our own experience in designing over 2,000 knock-ins for the OpenCell project (1), in which we systematically characterize the function of human genes by using fluorescent tags to measure the localization and interactions of the corresponding proteins. While we have implemented a sensible set of default design parameters based on the results presented in Figures 2-4, protoSpaceJAM is built to be fully customizable. The web application provides the user with a broad set of feature choices, which can be further tailored to specific applications by modifying the open-source design algorithm. By providing both the standalone open-source code and a user-friendly web interface at protospacejam.czbiohub.org, we aim to enable both expert and novice users to accelerate genetic research for a wide community.

### Limitations and future developments

In its current form, protoSpaceJAM is designed for HDR-based, insertional knock-in experiments using *S. pyogenes* Cas9 in three genomes (human, mouse, and zebrafish). The inclusion of additional genomes and Cas proteins (such as Cas12a, which has a TTTV PAM and is therefore better suited for AT-rich genomes) is an immediate direction for future development. Generalizing our algorithm to support knock-in applications for replacement mutagenesis would also broaden its impact and utility, for example to streamline the design of reagents for the introduction of disease-causing polymorphisms. Furthermore, while HDR-based strategies remain the most widely used for knock-in, the development of new approaches such as prime editing is rapidly changing the knock-in method landscape (46). Prime editing bypasses the requirement for double-strand breaks by using a Cas9 nickase fused to a reverse-transcriptase domain, together with a prime editing guide RNA (pegRNA) that bridges the Cas9 gRNA and the insertion payload, which can be integrated following reverse-transcription to its DNA equivalent (7). The design of prime editing experiments would therefore require a new set of constraints, which could be added to protoSpaceJAM.

Ultimately, the CRISPR field is characterized by constant innovation, with new methods for genome engineering being developed at a rapid pace. Bioinformatics tools for CRISPR design must therefore be adaptable. The power of open-source tools such as protoSpaceJAM, whose code is written in a modular format, lies in the ease of reuse and adaptation of code elements by anyone in the community, to pave the way for the development of ever-expanding toolsets to propel biological research.

## DATA AVAILABILITY

protoSpaceJAM and GenoPrimer are implemented in Python and are freely available as both a web server and standalone software under the BSD-3 license. The web server can be accessed at protospacejam.czbiohub.org/ without a login process. The standalone versions can be downloaded from the GitHub repository: github.com/czbiohub-sf/protoSpaceJAM and github.com/czbiohub-sf/GenoPrimer. The internal database of precomputed gRNA information can be downloaded following the usage instructions in the standalone versions.

## SUPPLEMENTARY DATA

Supplementary Data are available.

## AUTHOR CONTRIBUTIONS

Duo Peng: Conceptualization, Formal analysis, Methodology, Validation, Writing—original draft, review & editing. Madhuri Vangipuram: Formal analysis, Validation. Joan Wong: Conceptualization, Writing—review & editing. Manuel Leonetti: Conceptualization, Formal analysis, Methodology, Validation, Writing—original draft, review & editing.

## ACKNOWLEDGEMENTS

We thank J. Hanks, C. Kapfer, and R. Alvarez for help with computational resource support. We thank S. Schmid for critical feedback. We thank our donors for funding this work.

---

**Supplementary Figure 1.**
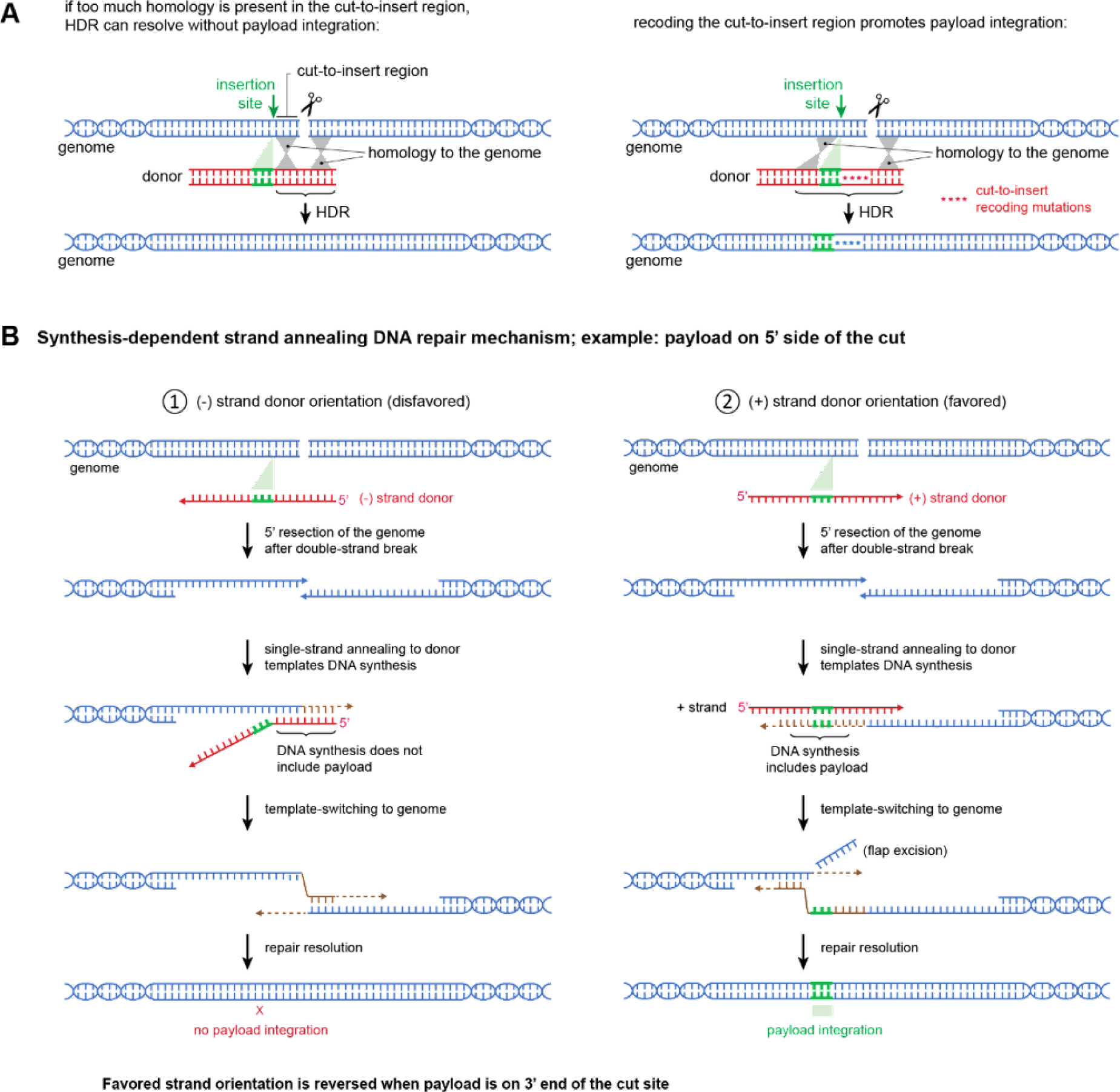
Recoding and strand selection impacts the rate of payload integration. **(A)** Schematic illustration of HDR resolving without payload integration due to homology between the target genome and the donor in the cut-to-insert region (left). “Repair track” mutations to reduce homology by recoding the cut-to-insert region promotes payload integration (right). **(B)** Schematic illustration of strand choice impacting payload integration as a function of cut site position relative to the insertion site with ssODN donors. ssODNs template DNA repair via a synthesis-dependent strand annealing mechanism. In the illustrated example, the insertion site is located on the 5’ side of the cut site. In this orientation, the plus strand (right) is more favorable for payload integration compared to the minus strand (left). The favored orientation will be reversed when the payload is on the 3’ side of the cut site.

**Supplementary Figure 2.**
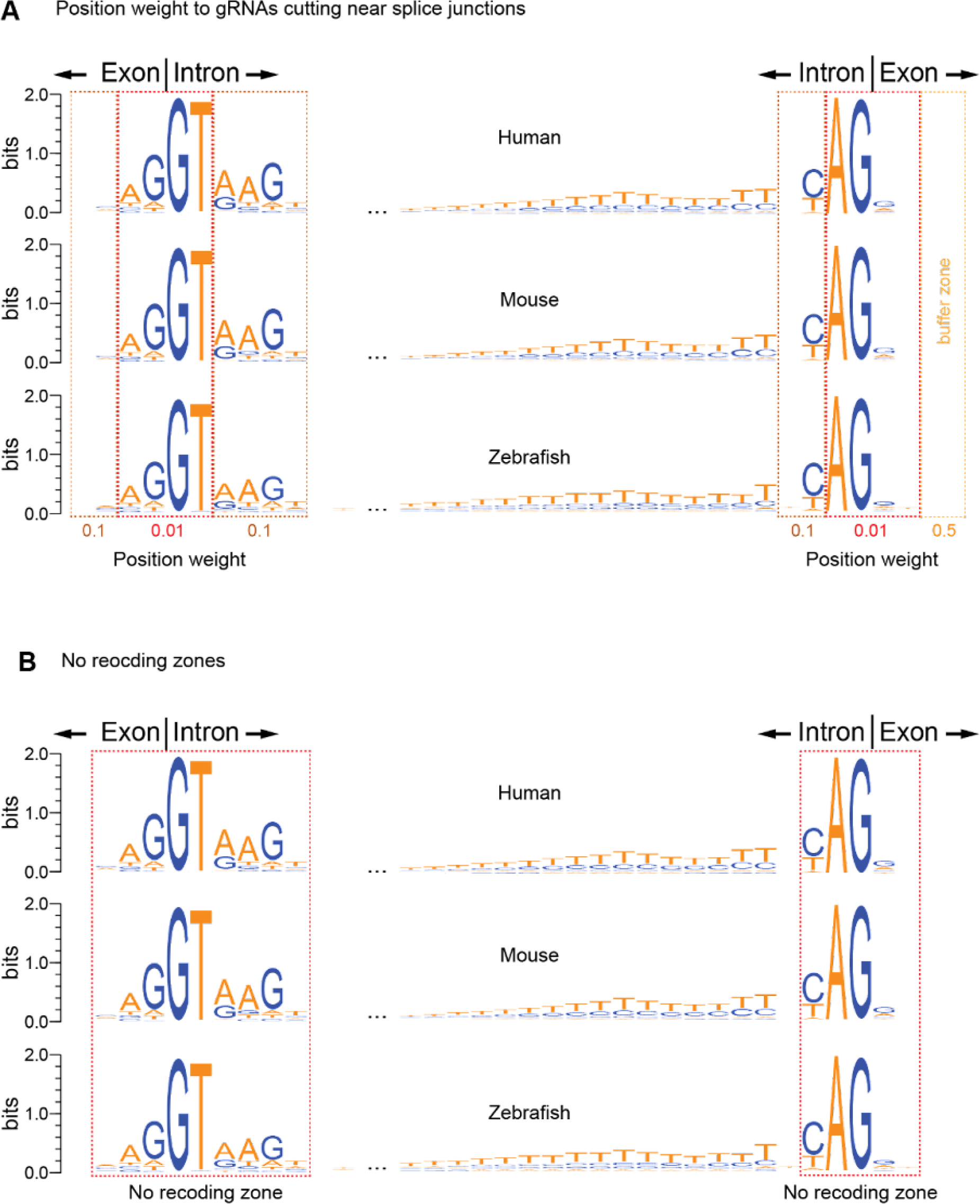
gRNA weights and recoding exclusion zones near splice junctions. (A-B) Sequence logo plots showing conservation of sequences near splice junctions in human, mouse, and zebrafish, using 1373460, 719549, and 848492 exon-intron and intron-exon junctions extracted from all annotated transcripts. **(A)** Different position weights are assigned to gRNAs according to the level of sequence conservation in different regions of the splice junctions. **(B)** Around exon-intron junctions, recoding is strictly prohibited within 3 bp upstream and 6 bp downstream of splice sites. Similarly, in the case of intron-exon junctions, recoding is prohibited within 3 bp upstream and 2 bp downstream of splice sites.

